# Selective targeting of the oligodendroglial GPR17 receptor improves myelin integrity and motor function in female SOD1^G93A^ mice

**DOI:** 10.64898/2026.04.28.721299

**Authors:** Stefano Raffaele, Tiziana Bonifacino, Francesca Carolina Mannella, Nhung Nguyen, Carola Torazza, Davide Marangon, Elena Maria Chinosi, Henrik Daa Schrøder, Eva Kildall Hejbøl, Kirsten Madsen, Laura Marchetti, Maria Letizia Trincavelli, Marco Milanese, Davide Lecca, Kate Lykke Lambertsen, Giambattista Bonanno, Maria P. Abbracchio, Marta Fumagalli

## Abstract

Amyotrophic lateral sclerosis (ALS) is a fatal neurodegenerative disease with no definitive disease-modifying therapies available, underscoring the urgent need to identify novel druggable targets. The G protein-coupled receptor GPR17 is a critical regulator of oligodendrocyte maturation and has emerged as a candidate target in ALS, yet its relevance to human disease and its therapeutic potential remain unclear. Here, we demonstrate that pathological GPR17 upregulation defines a conserved, pathologically immature oligodendroglial state in human ALS that can be pharmacologically leveraged to restore myelin integrity and improve functional outcome *in vivo*. Publicly available transcriptomics datasets and histological analyses revealed an increased abundance of GPR17-expressing immature oligodendrocytes in post-mortem human spinal cord tissue from ALS cases. Moreover, sustained activation of GPR17 with a selective agonist induced GPR17 internalization in heterologous expression systems and promoted the differentiation of primary oligodendrocyte precursors derived from SOD1^G93A^ mice. Translating these findings *in vivo*, chronic treatment with a brain-penetrant GPR17 agonist derived from the same pharmacological class significantly extended survival, delayed body weight loss, and improved motor performance in female SOD1^G93A^ mice, whereas male mice showed no therapeutic benefit. These effects were associated with restored oligodendrocyte maturation, preserved myelin integrity, motor neuron survival, and attenuated reactive gliosis in the spinal cord of female SOD1^G93A^ mice, while milder effects were observed in males. Together, these findings establish oligodendroglial GPR17 as a conserved and pharmacologically actionable target in ALS and show that sustained in vivo GPR17 agonism can reprogram altered oligodendroglial states and slow disease progression in a sex-dependent manner.

## 1. Introduction

Amyotrophic lateral sclerosis (ALS) is a relentlessly progressive neurodegenerative disease characterized by degeneration of upper and lower motor neurons, leading to muscle weakness, paralysis and death within a few years from diagnosis [1]. Despite substantial advances in understanding disease genetics and molecular pathology, effective pharmacological interventions remain scarce, highlighting the urgent need to identify novel, mechanistically grounded therapeutic targets [2]. Increasing evidence indicates that ALS progression is driven not only by intrinsic neuronal vulnerability but also by maladaptive interactions with non-neuronal cells, positioning glial populations as key modulators of disease evolution and attractive therapeutic targets [3].

Among glial cells, oligodendrocyte dysfunction has emerged as a critical yet underexploited contributor to ALS pathogenesis [4–6]. Oligodendrocytes are essential for axonal metabolic support, myelin maintenance, and long-range motor circuit integrity [4]. Analyses of post-mortem human tissue and studies in ALS models consistently reported early myelin abnormalities, defective oligodendrocyte turnover and impaired remyelination, which precede or accompany motor neuron degeneration [7,8]. Notably, these alterations are associated with an accumulation of oligodendrocyte precursor cells (OPCs) and immature oligodendrocytes, suggesting a failure of differentiation which contributes to motor deficits [7–9]. These observations raise the possibility that restoring oligodendrocyte maturation may represent a viable pharmacological strategy to slow ALS progression.

The G protein–coupled receptor GPR17 is a pivotal regulator of oligodendrocyte lineage differentiation and an attractive drug target [10,11]. GPR17 is transiently expressed during the transition from OPCs to immature oligodendrocytes, and its downregulation in immature oligodendrocytes is then required for full oligodendrocyte maturation and myelination [12–14]. Consistently, forced GPR17 overexpression locks cells in an immature state, impairing oligodendrocyte maturation and myelination, and reinforcing its role as a molecular brake on terminal differentiation [15,16]. Importantly, GPR17 belongs to the membrane rhodopsin-like GPCR family, a receptor class that historically has proven highly amenable to pharmacological modulation, underscoring its translational potential [10,11]. After the initial identification of uracil nucleotides and cysteinyl leukotrienes as endogenous ligands for GPR17 [17], more recent evidence has expanded this view by identifying also oxysterols as candidate endogenous modulators of GPR17 signaling [18,19]. This evolving pharmacology further supports the concept that GPR17 integrates extracellular metabolic and inflammatory cues to regulate oligodendrocyte maturation, exhibiting differential responses depending on specific pathophysiological contexts.

Previous work from our group has demonstrated a pathological upregulation of GPR17 in the spinal cord of the SOD1^G93A^ mouse model of ALS, where it marks an expanded population of immature oligodendrocytes with compromised differentiation capacity [9]. Notably, treatment with the marketed drug montelukast, a non-selective GPR17 antagonist, significantly counteracted the differentiation defects of SOD1^G93A^ OPCs *in vitro* and *in vivo*, restoring oligodendrocyte maturation and myelin integrity, preserving motor neurons, and improving survival and disease outcome of female SOD1^G93A^ mice [20]. Together, these findings suggest that reducing pathological GPR17 signaling may be beneficial in ALS. However, the relevance of GPR17 dysregulation to human ALS and the therapeutic value of selectively targeting GPR17 remain largely unexplored. In this context, selective GPR17 agonists may represent an alternative therapeutic strategy, as sustained stimulation can promote receptor desensitization, internalization and functional downregulation, thereby releasing the differentiation block associated with chronic GPR17 overexpression [16,21–24].

On this basis, in the present study, we aimed to establish the translational relevance of GPR17 as a pharmacological target in ALS and to assess the therapeutic impact of its selective modulation in the well-established SOD1^G93A^ mouse model. We first examined whether GPR17 dysregulation observed in ALS mouse models is conserved in human disease by integrating publicly available bulk and single-nucleus transcriptomic datasets from post-mortem spinal cord tissue with cell-type–resolved histological validation. We then investigated the effects of a selective GPR17 agonist on OPC differentiation *in vitro*, using primary cultures from the spinal cord of SOD1^G93A^ mice. Finally, we evaluated the *in vivo* efficacy of chronic treatment with galinex (GAL), a second-generation, brain-penetrant selective GPR17 agonist with optimized pharmacokinetic properties [24], in SOD1^G93A^ mice of both sexes.

Our findings demonstrate that GPR17 alterations are conserved in human ALS and that pharmacological modulation of GPR17 in experimental ALS models restores oligodendrocyte maturation and myelin integrity, confers neuroprotection, attenuates secondary gliosis, and improves motor performance, while revealing a marked sex-dependent therapeutic profile. Together, these results identify the oligodendroglial GPR17 receptor as a conserved and druggable target in ALS.

## 2. Materials and Methods

### 2.1 Bioinformatic analysis of publicly available datasets

To assess *GPR17* expression in human ALS spinal cord, we reanalyzed a published bulk RNA-seq dataset comprising post-mortem cervical, thoracic, and lumbar spinal cord sections from ALS cases and non-neurological disease controls [25]. Raw files were downloaded from Zenodo (https://zenodo.org/records/6385747) and used for downstream statistical analyses. Expression values are reported as log_2_TPM+1.

To estimate the abundance of *GPR17*-expressing cells, we queried a published single-nucleus RNA-seq dataset of post-mortem cervical spinal cord from 8 ALS cases and 4 non-neurological disease controls [26]. Data were explored via the Bioturing portal (https://talk2data.bioturing.com/sanofibrowser/) to extract representative UMAP plots of oligodendrocyte lineage cells and per-sample proportions of GPR17-expressing immature oligodendrocytes. For each sample, the number of GPR17 cells was computed and normalized to the total nuclei profiled for that sample, yielding a per-sample percentage that was subsequently normalized across replicate samples for comparative analyses.

### 2.2 Human tissue samples

Post-mortem human brain and lumbar spinal cord tissue was obtained from the Department of Pathology, Odense University Hospital (OUH, Denmark), in accordance with guidelines from the Health Ethics Committees for the Region of Southern Denmark (Journal number S-20220018). The cohort included 5 ALS cases (4 females, 1 male; median age = 74 years, IQR 69–75) and 7 non-neurological disease controls (2 females, 5 males; median age = 69 years, IQR 57–76). Age, sex, diagnosis, genetics, and disease duration are reported in Supplementary Table 1.

Formalin-fixed, paraffin-embedded (FFPE) samples were cut into 2 μm sections on a microtome and processed for immunohistochemistry as previously described [27,28].

### 2.3 Immunohistochemistry and image analysis on human tissue

Immunofluorescence and brightfield immunohistochemistry were performed on FFPE sections following previously established protocols [27]. Briefly, sections were pretreated with Autofluorescence Eliminator Reagent (Merck Millipore, Merck KGaA, Darmstadt, Germany) and incubated overnight at 4 °C with the following primary antibodies: rabbit anti-GPR17 (1:300; custom-made by PRIMM, Milan, Italy), mouse anti-IBA1 (1:1,000; Sigma-Aldrich, Merck KGaA, Søborg, Denmark), mouse anti-GFAP-Cy3 (1:500; Sigma-Aldrich, Merck KGaA), mouse anti-CNPase (1:100; Santa Cruz Biotechnology Inc., Dallas, TX, USA), mouse anti-BCAS1 (1:500, Santa Cruz Biotechnology Inc.), and mouse anti-NeuN (1:100, Merck Millipore). Appropriate fluorophore-conjugated secondary antibodies were applied the following day: goat anti-rabbit Alexa Fluor 594, goat anti-rabbit Alexa Fluor 488, or goat anti-mouse Alexa Fluor 488 (all 1:400, Thermo Fisher Scientific Inc., Waltham, MA, USA), and nuclei were counterstained with 4’,6-diamidine-2’-phenylindole dihydrochloride (DAPI; 1:1,000; Thermo Fisher Scientific Inc.). For chromogenic detection of GPR17 (rabbit anti-GPR17, custom PRIMM), sections were incubated with Dako EnVision®+Dual Link System-HRP (Dako, Agilent Technologies Denmark ApS, Glostrup, Denmark) followed by 3,3’-diaminobenzidine (DAB) solution.

Images were acquired at 40× magnification with a Nikon ECLIPSE Ti2 confocal microscope (Nikon, Tokyo, Japan) for immunofluorescence and with a NanoZoomer slide scanner (Hamamatsu Photonics K.K., Shizouka, Japan) for immunohistochemistry. Quantitative analyses were performed using NDP.view2 software (Hamamatsu Photonics K.K.), with manual counting of GPR17-positive cells by an investigator blinded to experimental groups. Only cells with a visible haematoxylin-stained nucleus were included. Cell density was expressed as the number of positive cells per mm^2^, normalized to the analyzed tissue area.

### 2.4 Ligand-induced GPR17 cellular internalization assay

1321N1 human astrocytoma cells, lacking endogenous expression of P2Y and CysLT receptors including GPR17 [17], were maintained in complete DMEM/F12 medium (Corning, New York, USA) supplemented with 10% fetal bovine serum (FBS, Corning, New York, USA) and penicillin–streptomycin (1%, Corning, New York, USA) at 37°C in 5% CO2. Cells were seeded at a density of 50,000 cells per well in 24 well plates, each containing a previously sterilized coverslip, and were subsequently transfected with a plasmid encoding a GFP-tagged GPR17 fusion protein (pEGFPN1–GPR17, Clontech) using JetPEI® transfection reagent (PolyPlus). After 48 h, cells were treated in complete medium with either Asinex 1 (ASN, 10 nM), LTD4 (10 µM), 22(R)-hydroxycholesterol (OXY, 1 µM), or Cangrelor (CGL, 10 µM) for 30 or 60 min. Control samples were exposed to vehicle (0.1% DMSO or 0.1% ethanol) for the corresponding durations. Following treatment, cells were first fixed with a 4% paraformaldehyde solution in phosphate-buffered saline (PBS), then mounted with Fluoroshield Mounting Medium (F6057-20 ML-Sigma) on a microscope support. Samples were imaged with a laser scanning confocal microscope (Nikon Eclipse Ti-A1 MP) using a 60x/1.4NA oil objective and pinhole set to 1 Airy Unit. To ensure analysis within an intracellular optical section, images were acquired at the focal plane in which the nucleus appeared sharply focused. Under these conditions, the main plasma membrane pool was detected as the cell perimeter, allowing discrimination between intracellular and membrane-associated fluorescence signals. The 405 and 488 laser lines were used for imaging DAPI and GFP by using the 400−500 and 475−575 nm emission filter cubes, respectively. Acquisition parameters were maintained constant throughout the analysis. All images were analysed by ImageJ program. For each field, cells were categorized based on receptor subcellular localization in three sub-populations (membrane: when most GPR17 signal stems from plasma membrane, i.e. cell perimeter; membrane plus intracellular vesicles: where both membrane and vesicle signal is visible; internalized: where almost no membrane signal is appreciable and the receptor signal was mainly intracellular).

### 2.5 Animal model

Transgenic SOD1^G93A^ mice (B6SJL-TgN SOD1/G93A1Gur; [29]) were used, as previously described [20,30]. This line, which expresses a high copy number of mutant human SOD1 with a Gly93Ala substitution, is the most widely used ALS preclinical model and reproduces key pathological features of the human disease [31]. The colony was maintained by crossing SOD1^G93A^ males with background-matched B6SJL wild-type (WT) females, thereby keeping the transgene in the hemizygous state. Genotyping was performed as previously reported in [32]. Mice develop clear ALS-like symptoms at approximately 12–13 weeks of age and reach the late symptomatic stage around 16–18 weeks. Animals were housed six to seven per standard cage (42 × 27 × 16 cm; floor surface 1.134 cm²) under controlled conditions (temperature 22 ± 1 °C; humidity 50%) with a 12-h light/dark cycle (lights on 07:00–19:00). Standard chow diet (4RF21 diet; Mucedola, Settimo Milanese, Milan, Italy) and water were available ad libitum. Environmental enrichment (plastic igloos or paper rolls) was provided and regularly changed. Animals were monitored daily to assess health status. Both sexes were included in the study. All procedures were conducted in compliance with the European Directive 2010/63/EU for the protection of animals used for scientific purposes, the Italian Legislative Decree No. 26/2014, and the recommendations of the ARRIVE guidelines [33]. The experimental protocol was approved by the Italian Ministry of Health (Project Authorization No. 97/2017-PR and 1022/2020-PR). Survival studies were terminated before spontaneous death, following established clinical scores to individuate the humane endpoint, corresponding to complete hindlimb paralysis combined with loss of the righting reflex [34,35]. For animals included in the histological analyses, sacrifice was performed at postnatal day 125 (P125), corresponding to the late symptomatic stage.

### 2.6 Primary OPC culture experiments

Primary OPCs were isolated from spinal cords of P7 SOD1^G93A^ mice, as previously described [9]. Spinal cords were dissected at 4 °C and maintained in Tissue Storage Solution (Miltenyi Biotec) until dissociation. Tissue was enzymatically dissociated using a papain-based Neural Tissue Dissociation Kit (Miltenyi Biotec), and PDGFRα OPCs were purified by magnetic-activated cell sorting (MACS) following incubation with anti-PDGFRα microbeads (Miltenyi Biotec), according to the manufacturer’s instructions. Approximately 60,000 OPCs were obtained per pup and plated onto poly-D-ornithine–coated 24-well plates at a density of 30,000 cells/well. Cells were maintained in OPC proliferation medium consisting of Neurobasal medium supplemented with 2% B27, 1% L-glutamine, 1% penicillin/streptomycin, 10 ng/mL PDGF-AA and 10 ng/mL FGF2. After 2 days *in vitro*, cultures were switched to a differentiation medium containing DMEM supplemented with 1% N2, 2% B27, 0.01% BSA, 1% L-glutamine, 1% penicillin/streptomycin and 10 ng/mL triiodothyronine (T3). After 24 h in differentiation medium, cells were treated with the selective GPR17 agonist Asinex 1 (ASN; 1 µM) or an equivalent concentration of dimethyl sulfoxide (DMSO, vehicle control), and treatments were maintained for additional 48 h. Cells were then fixed in 4% paraformaldehyde and processed for immunocytochemistry as previously published [9], using rat anti-myelin basic protein (MBP, 1:200; cat. no. MAB386, Millipore, Milan, Italy) primary antibody followed goat anti-rat secondary antibody conjugated to Alexa Fluor 555 (1:600; Life Technologies). Nuclei were labelled with Hoechst33258. Fluorescent images were acquired using an inverted Zeiss 200M microscope, and quantitative analyses were performed in ImageJ on 20–40 randomly selected fields per coverslip (≥3 coverslips per condition). To assess brain-derived neurotrophic factor (BDNF) release, before fixation culture supernatants were collected, centrifuged to remove debris, and stored at −80 °C until analysis. BDNF levels were quantified using a commercially available ELISA kit (cat. No. EA100207, OriGene Technologies, Herford, Germany), following the manufacturer’s instructions.

### 2.7 *In vivo* pharmacological treatment

Pharmacological profiling of the selective GPR17 agonist galinex (GAL; [24]) has demonstrated high potency at the GPR17 receptor *in vitro* (EC = 0.64 ± 0.12 nM in a [³ S]GTPγS assay), together with favorable in vivo pharmacokinetic properties [24]. Importantly, brain drug metabolism and pharmacokinetics (DMPK) analysis following subcutaneous administration of GAL at 10 mg/kg showed robust CNS exposure (C_max = 270 ng/ml; T_max = 0.25 h; C_last = 5.80 ng/ml at T_last = 6.00 h; AUC_last = 205 h·ng/ml), indicating efficient brain penetration compatible with pharmacological engagement of GPR17 at the administered dose [24].

GAL (Amb9128576; Ambinter-Greenpharma, Orléans, France) was dissolved in a vehicle consisting of 50% DMSO and 50% PEG400 and delivered at a dose of 10 mg/kg/day by continuous subcutaneous infusion using Alzet osmotic minipumps (100 μl reservoir; infusion rate 0.11 μl/h; Cupertino, CA, USA). Minipumps were implanted at postnatal day 90 (P90), corresponding to the early symptomatic stage of disease progression in SOD1^G93A^ mice, thereby modelling a therapeutic rather than preventive treatment paradigm. Continuous drug delivery was maintained for at least 28 days. Control SOD1^G93A^ mice received vehicle (VEH) using the same surgical and infusion procedures.

### 2.8 Behavioral tests

The effects of GAL treatment on disease progression were evaluated by behavioral studies performed three times per week, starting at P80 (10 days before starting the treatment), until sacrifice. A total of n = 23 SOD1^G93A^ mice (12 females and 11 males) treated with GAL and n = 21 SOD1^G93A^ mice (10 females and 11 males) treated with VEH were included. Sample sizes were predetermined according to published recommendations for ALS preclinical studies [36]. Behavioral assessments were carried out in a blinded fashion, as previously described [20,35].

#### 2.8.1 Survival probability, body weight loss, and food intake

Survival was monitored from treatment onset until animals reached the humane endpoint, defined by complete hindlimb paralysis combined with loss of the righting reflex [34]. At this stage, mice were euthanized according to approved protocols. Body weight was recorded before each behavioral session as a marker of disease progression, whereas food intake was monitored every other day from P98 to P120, as allowed by animal conditions.

#### 2.8.2 Rotarod

Motor coordination was assessed using an accelerating rotarod apparatus (RotaRod 7650, Ugo Basile, Gemonio, VA, Italy). The rotation speed increased from 4 to 40 rpm over 5 min, and latency to fall was recorded. Mice underwent a 1-week training period prior to testing.

#### 2.8.3 Beam balance

Balance was tested on a 1-m-long, 6-mm-wide beam elevated 50 cm above the ground. Animals were encouraged to cross the beam toward a dark box positioned at the end, and the number of foot slips was counted [37].

#### 2.8.4 Extension reflex

Neuromuscular deficits were evaluated by observing hindlimb posture while suspending mice by the tail. Reflexes were scored on a 5-point scale ranging from 5 (full extension) to 0 (complete impairment), as described previously [30,35].

#### 2.8.5 Gait impairment

Gait was assessed by direct observation of spontaneous locomotion in an open field. A 5-point scoring system was used, with higher scores corresponding to preserved motor ability [30,35].

#### 2.8.6 Grip Strength

Forelimb muscle strength was measured using a GSM Grip-Strength Meter (Ugo Basile, Gemonio, VA, Italy), equipped with a force transducer (capacity 1500 g, resolution 0.1 g). Mice were gently suspended by the tail and allowed to grasp the horizontal pull bar connected to the dynamometer. Once the grip was established, animals were steadily pulled backwards by the tail until they released the bar, and the maximal opposing force was automatically recorded. Each trial consisted of at least three consecutive measurements, and mean values were calculated after excluding outliers.

#### 2.8.7 Hanging wire

Hindlimb endurance was evaluated by placing mice on a wire grid suspended 50 cm above bedding, then inverting the grid. The latency to release the grid with both hindlimbs was recorded, with a cut-off of 120 s.

### 2.9 Immunohistochemistry on rodent tissue

Mice used for immunohistochemical analyses were not enrolled in behavioral studies. To reduce animal use, spinal cord tissue from the same animals was employed for all immunostaining experiments. Two groups of SOD1^G93A^ mice were analyzed: GAL-treated (10 mg/kg/day) and VEH-treated (n = 10 per group, 5 females and 5 males). At the late symptomatic stage (P125), animals were deeply anesthetized and transcardially perfused with PBS. Spinal cords were dissected, fixed in 4% paraformaldehyde, cryoprotected in 30% sucrose, embedded in optimal cutting temperature compound (OCT), and cut into 20-µm sections using a cryostat, as already described [20]. Immunofluorescence protocols followed previously established procedures [9,20,38,39]. Briefly, sections were blocked and incubated with the following primary antibodies: rabbit anti-NG2 (1:2,000; Cat. No. AB5320, Merck, Milan, Italy), rabbit anti-GPR17 (1:2,500; custom-made by PRIMM, Milan, Italy; [9,40]), rat anti-MBP (1:200; Cat. No. MAB386, Merck, Milan, Italy), rabbit anti-Iba1 (1:500; Cat. No. 019–19741, Wako, Osaka, Japan), mouse anti-glial fibrillary acidic protein (GFAP, 1:300; Cat. No. 3670, Cell Signaling Technologies, Leiden, The Netherlands), goat anti-Serpina3n (1:100, Cat. No. AF4709, R&D Systems, Abingdon, United Kingdom), rabbit anti-aspartoacylase (ASPA, 1:100; ABN1698, Merck; Milan, Italy), rabbit anti-Hb9 (1:500; Cat No. ABN174, Merck, Milan, Italy), and mouse anti-neurofilament heavy chain (NF-H) clone SMI32 (1:500; Cat. No. 801701, BioLegend, Amsterdam, The Netherlands), rabbit anti-OLIG2 (1:600; Cat. No. AB9610, Merck, Milan, Italy). Heat-induced epitope retrieval was performed for GPR17 staining with 10 mM of citrate buffer pH 6 (Merck, Milan, Italy) containing 0.05% Tween20 (Merck, Milan, Italy). Appropriate secondary antibodies were used: goat anti-rabbit Alexa Fluor 555 (Cat. No. A21428), goat anti-rabbit Alexa Fluor 488 (Cat. No. A11008), goat anti-mouse Alexa Fluor 555 (Cat. No. A21422), goat anti-rat Alexa Fluor 488 (Cat. No. A11006), and donkey anti-goat Alexa Fluor 555 (Cat. No. A21432) (1:600; all from Life Technologies, Monza, Italy). Signal amplification was performed for GPR17 and NG2 staining, using High Sensitivity Tyramide-Rhodaminate Signal Amplification kit (Akoya Biosciences, Monza, Italy), following the manufacturer’s instructions. Nuclei were counterstained with Hoechst 33258 (0.3 μg×ml-1; Life Technologies, Monza, Italy). All staining steps, imaging, and subsequent analyses were performed in parallel for GAL- and VEH-treated groups under identical conditions.

### 2.10 Image acquisition and quantitative analysis

Images were acquired from the ventral gray matter of the lumbar spinal cord, with two to three sections analyzed per animal at 40× magnification using a Nikon ECLIPSE Ti2 confocal microscope (Nikon, Tokyo, Japan). All analyses were performed by an investigator blinded to the experimental groups using Fiji/ImageJ software. Identical acquisition settings and analysis procedures were applied across GAL- and VEH-treated groups.

Quantitative evaluations included manual cell counting with the “Cell Counter” tool, normalized to the total analyzed area to obtain the density (cells/mm²) of NG2-, GPR17-, ASPA-, and Hb9-positive cells. In addition, the positive area fraction was measured for Iba1, GFAP, Serpina3n, and MBP immunostainings.

Myelin integrity and the proportion of myelinated axons in the region of interest were quantified in Fiji/ImageJ using the Color Threshold plugin. Lumbar spinal cord sections were double-immunolabeled for MBP (myelin) and NF-H (axons). Images were converted to 24-bit RGB (8-bit/channel) and thresholded to separate specific signal from background, generating binary masks for MBP, NF-H, and their co-localization. Positive area fractions were quantified, and a myelination index was calculated as the MBP /NF-H co-localization area normalized to total NF-H area.

### 2.11 Spectral confocal reflectance microscopy of myelin integrity

Spectral confocal reflectance (SCoRe) microscopy was used to visualize compact CNS myelin based on its higher refractive index relative to the surrounding parenchyma, enabling detection of individual myelin internodes by reflected light [41,42]. Reflectance images were acquired on a Nikon AX R confocal microscope equipped with a 20/80 dichroic beam splitter, using excitation at 488, 561 and 640 nm and three photodetectors with narrow detection bandwidths centred on each laser line. Laser power and detector gain were initially set to minimum and progressively increased until the first saturated pixel appeared, then slightly reduced to eliminate saturation; parameters were determined on the first image and kept constant thereafter. Ventral gray matter images were collected from 20-µm-thick lumbar spinal cord sections mounted in Dako fluorescence mounting medium, using a 40× water-immersion objective. For quantification, reflectance images from each laser line were merged into a single composite image and a minimum intensity threshold was applied in Fiji/ImageJ to identify compact myelin-positive pixels; the SCoRe signal was expressed as compact myelin area fraction.

### 2.12 Statistical analysis

Statistical analyses were performed using Prism 10 software (GraphPad, San Diego, CA, USA). Data are expressed as mean ± standard error of the mean (SEM). Normality of distributions was assessed using the Shapiro–Wilk test for sample sizes < 8. Survival probability was analyzed using Kaplan–Meier plots and compared with the log-rank test. Comparisons between two groups were performed using unpaired Student’s t-test for normally distributed data or Mann–Whitney U test for non-parametric data. Repeated measures from behavioral studies were analyzed with mixed-effects models. Outliers were identified and excluded using the Robust Regression and Outlier Removal (ROUT) method (Q = 1). Statistical significance was set at p < 0.05. Specific details for each analysis are reported in the corresponding figure legends.

## 3. Results

### 3.1. Identification of the oligodendroglial GPR17 receptor as a conserved pharmacological target in human ALS

Considering our previous work demonstrating increased GPR17 protein levels and accumulation of GPR17-expressing immature oligodendrocytes in the lumbar spinal cord of the SOD1^G93A^ mouse model of ALS [9,20], we investigated whether similar alterations are present in human disease. Reanalysis of a published bulk RNA-sequencing dataset from post-mortem human spinal cord samples derived from patients carrying distinct ALS-linked mutations (including *C9orf72, TARDBP, SOD1, FUS, ANG, OPTN,* and sporadic cases with unknown genetic association; [25]) revealed a significant upregulation of *GPR17* transcript in ALS cases compared with non-neurological controls (CTRL), consistently across cervical, thoracic and lumbar segments (Figure 1A). To determine whether this increase reflected changes in specific cellular populations, we interrogated an independent single-nucleus RNA-sequencing dataset of human cervical spinal cord [26], which demonstrated a selective increase in the proportion of GPR17-expressing immature oligodendrocytes in sporadic ALS samples compared to CTRL, indicating an expansion of this lineage state in the diseased spinal cord (Figure 1B-C).

**Figure 1.**
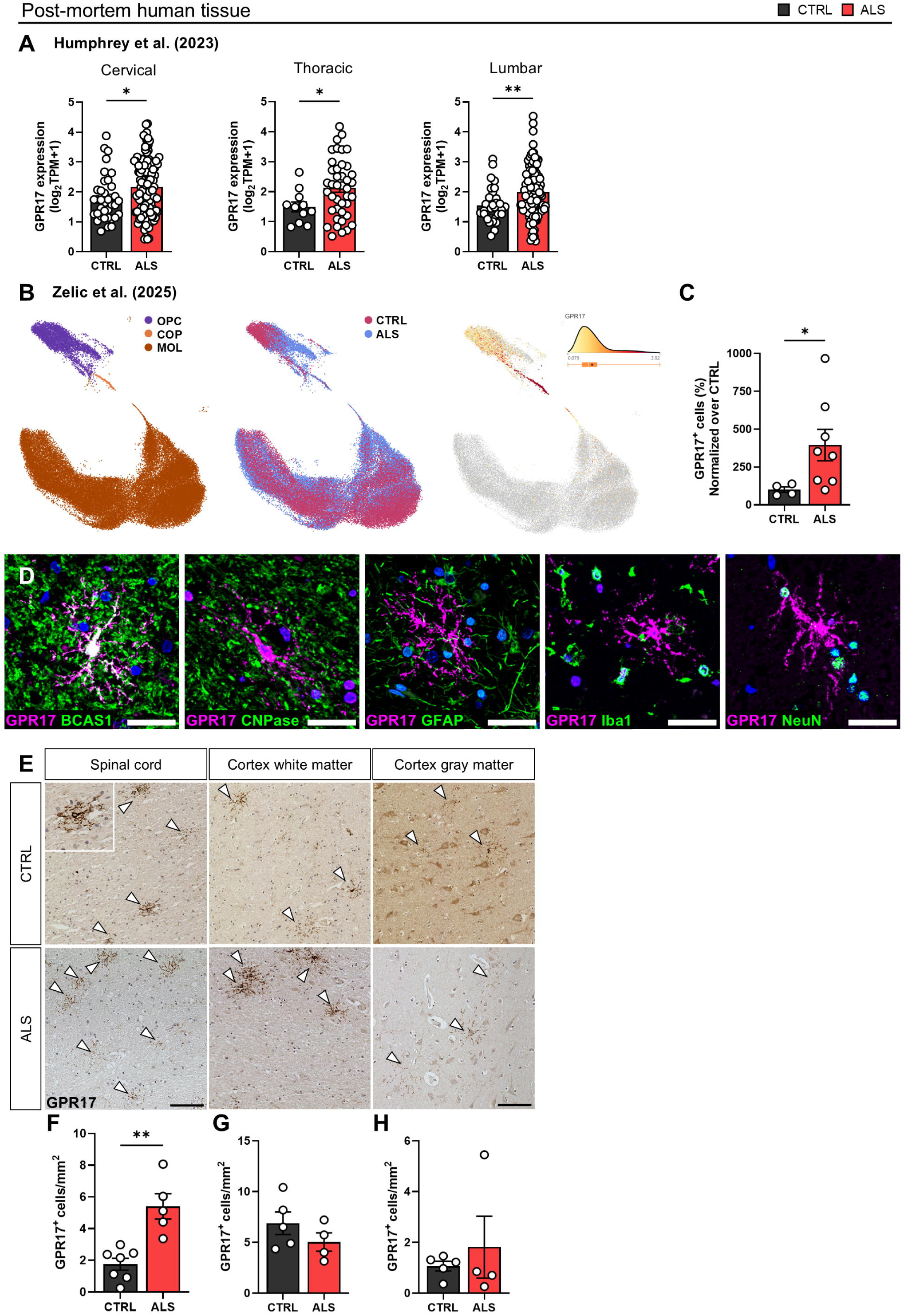
Analysis of GPR17-expressing immature oligodendrocytes in human ALS tissue. (**A**) GPR17 expression in post-mortem human spinal cord from ALS cases and non-neurological disease controls (CTRL), obtained from published bulk RNA-seq datasets [25]. Cervical (CTRL: n = 36; ALS: n = 138), thoracic (CTRL: n = 11; ALS: n = 41) and lumbar (CTRL: n = 33; ALS: n = 119) segments are shown. *p < 0.05, **p < 0.01, Mann–Whitney test. (**B**) UMAP plots of oligodendrocyte lineage cells in post-mortem human cervical spinal cord tissue from ALS cases and CTRL, retrieved from a published single-nucleus RNA-seq dataset [26], color-coded by subcluster (OPC, COP, MOL), disease condition and GPR17 expression level. (**C**) Proportion of GPR17-expressing cells in post-mortem human cervical spinal cord tissue from ALS cases (n = 8) and CTRL (n = 4), derived from single-nucleus RNA-seq data [26]. *p < 0.05, Welch’s t-test. (**D**) Representative immunofluorescence images of GPR17 (magenta) co-localization with the immature oligodendrocyte marker BCAS1, but not with the mature oligodendrocyte marker CNPase, the astrocyte marker GFAP, the microglia/macrophage marker Iba1 or the neuronal marker NeuN (green), in post-mortem human lumbar spinal cord tissue. Nuclei are counterstained with DAPI. Scale bar: 100 μm. (**E**) Representative images of GPR17 cells (arrowheads) in post-mortem human lumbar spinal cord and cortical tissue from ALS cases and CTRL. Scale bar: 100 μm. (**F-H**) Quantification of GPR17 cell density in post-mortem human lumbar spinal cord (F), cortical white matter (G), and cortical gray matter (H) tissue from ALS cases and CTRL (n = 4-7). **p < 0.01, Student’s t-test.

At the protein level, immunohistochemical analyses of human post-mortem spinal cord and cortical samples from ALS cases (four sporadic and one carrying SOD1 mutation) and CTRL were performed to univocally define the cellular identity and density of GPR17 cells. Double-labelling experiments showed robust co-localization of GPR17 with the immature oligodendrocyte marker BCAS1, whereas no overlap was observed with the mature oligodendrocyte marker CNPase (Figure 1D). In addition, GPR17 immunoreactivity did not co-localize with markers of astrocytes (GFAP), microglia/macrophages (Iba1), or neurons (NeuN), confirming the selective association of GPR17 with immature oligodendroglial cells in human spinal cord tissue (Figure 1D). Subsequent quantitative analyses revealed a significant increase in the density of GPR17 cells in the spinal cord of ALS patients compared to CTRL (Figure 1E-F), while no differences were detected in cortical tissue (Figure 1E-H).

Collectively, these transcriptomic and histological data are consistent with a spinal cord– restricted expansion of GPR17-expressing immature oligodendrocytes in human ALS, closely mirroring observations in experimental models. The consistency of GPR17 dysregulation across species and analytical platforms supports its pharmacological relevance and highlights GPR17 as a promising target for therapeutic intervention in ALS.

### 3.2. Treatment with a selective GPR17 agonist fosters SOD1^G93A^ OPC differentiation *in vitro*

Building on our previous evidence that primary OPCs isolated from SOD1^G93A^ mice exhibit an intrinsic impairment in terminal differentiation associated with pathological GPR17 upregulation [9], we next investigated whether pharmacological modulation of GPR17 signaling could restore OPC maturation and functional output *in vitro*. Treatment of OPC cultures derived from SOD1^G93A^ mice with the selective GPR17 agonist asinex 1 (ASN, [23,43]) resulted in a significant enhancement of OPC differentiation, as shown by an increased proportion of mature MBP cells compared with vehicle-treated cultures (Figure 2A-B). In parallel, ASN treatment induced a significant increase in BDNF levels in the culture medium (Figure 2C), indicating that GPR17 engagement does not only promote oligodendroglial maturation but also fosters the trophic support provided by differentiating OPCs.

**Figure 2.**
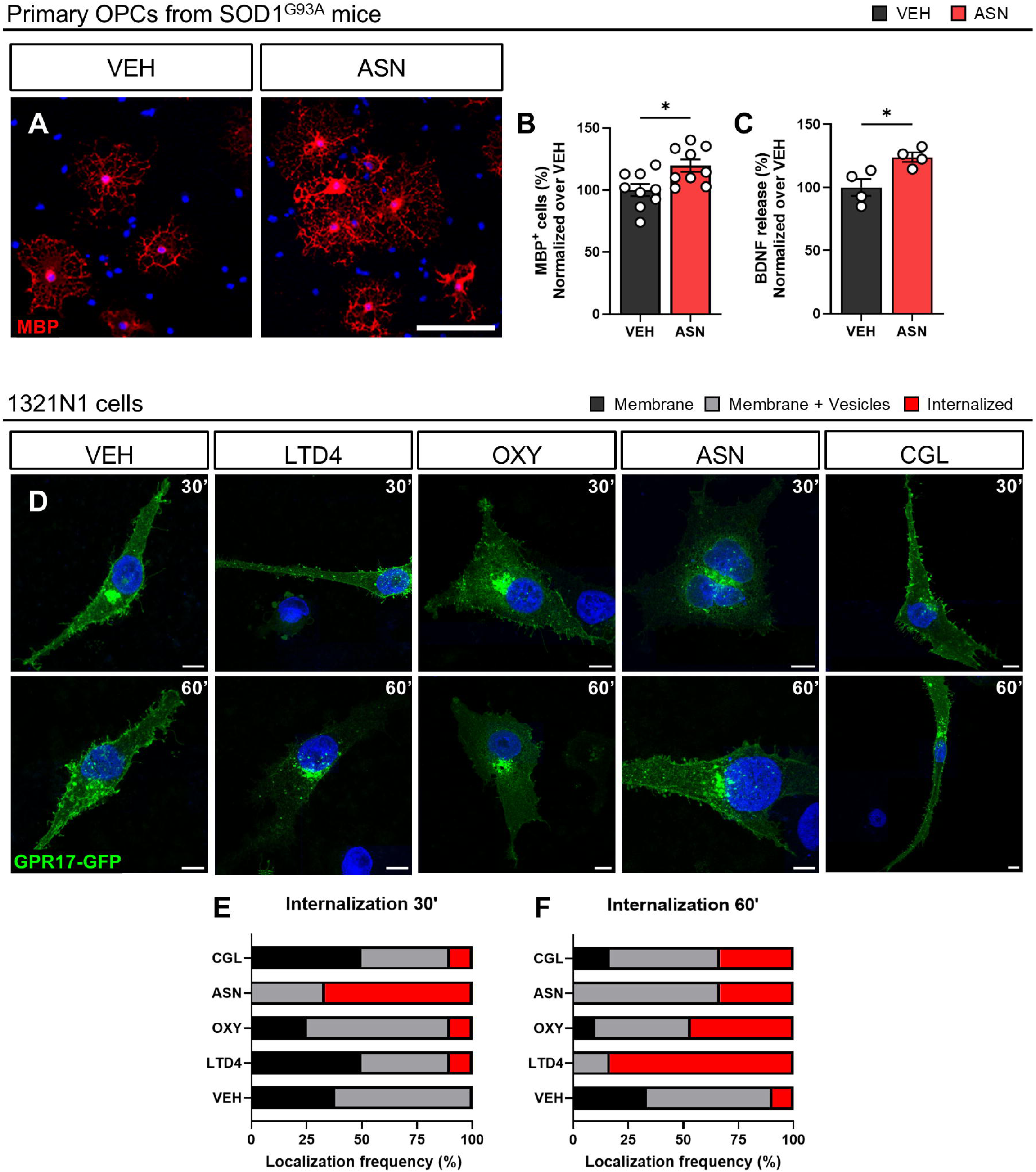
GPR17 internalization and maturation of OPCs isolated from the spinal cord of SOD1^G93A^ mice upon ASN treatment. (**A**) Representative images showing MBP^+^ mature cells in SOD1^G93A^ cultures exposed to ASN or VEH. Hoechst33258 was used to label cell nuclei. Scale bar: 100 μm. (**B**) Quantification of the percentage of MBP^+^ cells in SOD1^G93A^ cultures exposed to ASN or VEH (n = 9 coverslips from three independent experiments). * p < 0.05, Student’s t-test. (**C**) Quantification of BDNF protein levels in the medium of SOD1^G93A^ cultures exposed to ASN or VEH (n = 4 samples from two independent experiments). * p < 0.05, Student’s t-test. (**D**) Representative images showing 1321N1 cells (nuclei visible in blue channel) transfected with GPR17-GFP reporter (green channel) and treated with VEH, LTD4, ASN, or CGL for 30’ or 60’ during GPR17 internalization assay. Scale bar: 10 μm. (**E-F**) Quantification of the percentage of 1321N1 cells displaying GPR17 localization in the plasma membrane, both membrane and vesicles, or intracellular vesicles after treatment with VEH, LTD4, ASN, or CGL for 30’ (E) or 60’ (F; n = 5-7 technical replicates from two independent experiments).

To evaluate whether this pro-differentiative effect was mediated by agonist-induced receptor internalization [16,21,22], we established a reporter assay in 1321N1 cells transfected with a GFP-tagged GPR17. Relative to vehicle-treated controls, ASN induced a marked redistribution of GPR17 from the plasma membrane toward intracellular vesicular compartments, displaying a faster kinetic profile than the endogenous ligands LTD4 and OXY (Figure 2D-F). Conversely, the antagonist cangrelor (CGL) did not trigger internalization, remaining comparable to basal conditions (Figure 2D-F).

Hence, these findings demonstrate that, by promoting fast and efficient GPR17 internalization and removal from the plasma membrane, ASN rescues disease-associated differentiation defects in SOD1^G93A^ OPCs and enhances neurotrophic factor release, supporting GPR17 modulation as a potential strategy to improve oligodendroglial function and promote neuroprotection in ALS.

### 3.3. Chronic *in vivo* treatment with a selective GPR17 agonist extends survival rate and delays body weight loss in female SOD1^G93A^ mice

To translate our in vitro findings to an in vivo setting, we next evaluated the therapeutic impact of galinex (GAL), a second-generation GPR17 agonist derived from ASN through an iterative drug discovery optimization pipeline and characterized by improved pharmacokinetic properties for the *in vivo* application [24]. GAL was administered chronically to SOD1^G93A^ mice at the dose of 10 mg/kg/day via subcutaneous infusion with osmotic minipumps, starting at the early symptomatic stage P90. Chronic GAL treatment resulted in a significant extension of survival in female SOD1^G93A^ mice compared to vehicle-treated controls, whereas no survival benefit was observed in male littermates (Figure 3A-B). Consistent with this sex-specific effect on lifespan, GAL administration also significantly delayed body weight loss in female mice, a robust indicator of slower disease progression in this model, while body weight trajectories in male mice were unaffected (Figure 3C-D). Interestingly, body weight improvement was paralleled by increased food intake in GAL-treated female mice, whereas feeding behavior was not modified by the treatment in males (Figure 3E-F).

**Figure 3.**
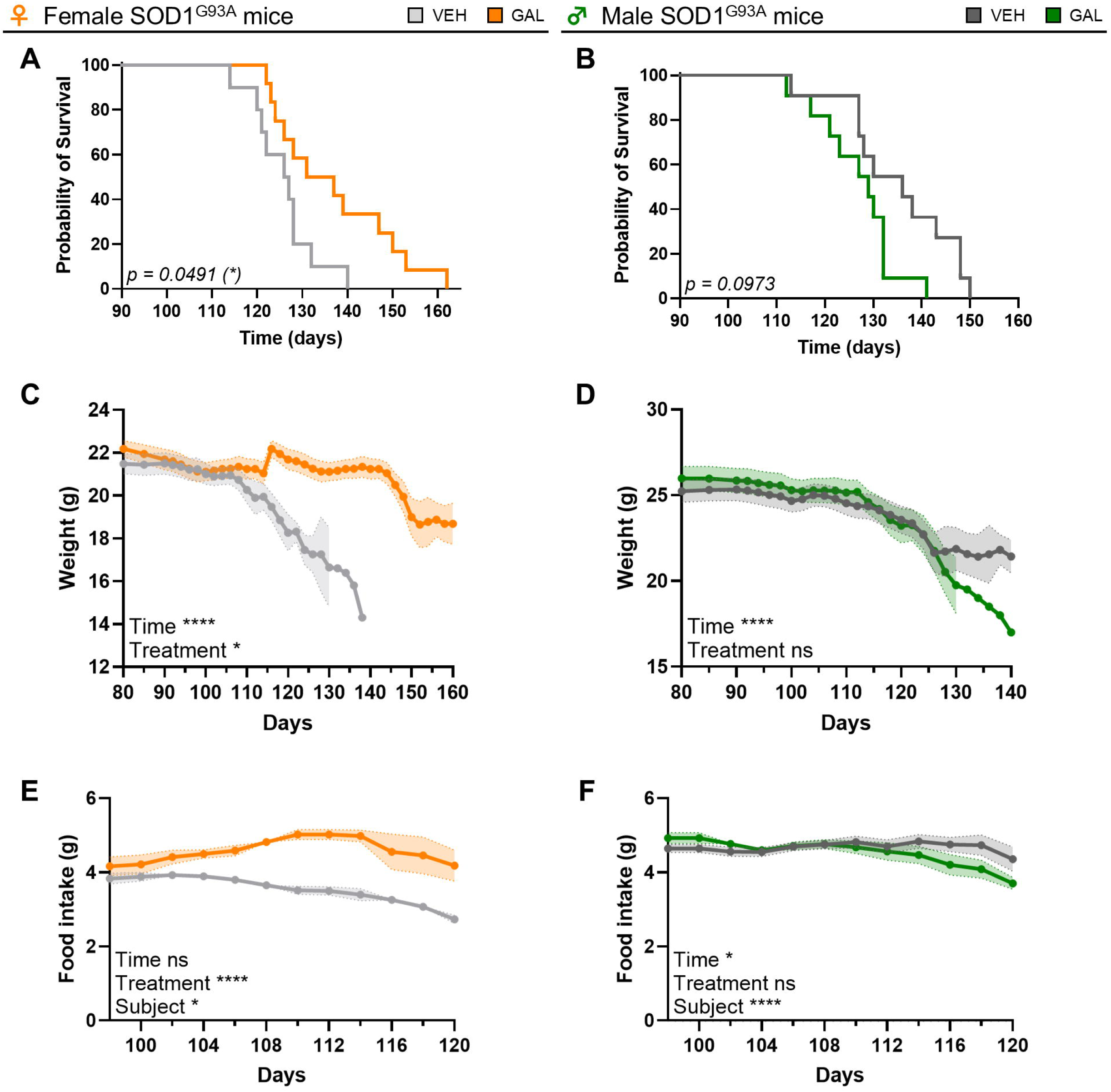
Survival probability, body weight loss, and food intake in GAL and vehicle treated SOD1^G93A^ mice. (**A-B**) Survival probability of female (A) and male (B) SOD1^G93A^ mice treated with vehicle (VEH) or 10 mg/kg/day GAL (n=10-12). * p<0.05, Kaplan–Meier analysis with Gehan-Breslow-Wilcoxon test. (**C-D**) Quantification of the body weight of female (C) and male (D) SOD1^G93A^ mice treated with VEH or 10 mg/kg/day GAL (n=10-12). * p<0.05, **** p<0.0001, mixed-effects model with Geisser-Greenhouse correction for repeated measures. (**E-F**) Quantification of food intake of female (E) and male (F) SOD1^G93A^ mice treated with VEH or 10 mg/kg/day GAL (n=6). * p<0.05, **** p<0.0001, two-way RM ANOVA.

Together, these data demonstrate that chronic pharmacological modulation of GPR17 with GAL exerts beneficial effects *in vivo* in a sex-dependent manner, selectively improving survival and delaying functional decline in female SOD1^G93A^ mice.

### 3.4. Chronic *in vivo* treatment with a selective GPR17 agonist ameliorates motor functions of female SOD1^G93A^ mice

We next compared the motor performance of GAL- and vehicle-treated SOD1^G93A^ mice, matched for age and sex. As to motor coordination, we observed a significant amelioration of rotarod performance and beam balance score in GAL-treated female SOD1^G93A^ mice versus vehicle, whereas GAL treatment was found to worsen rotarod performance in male SOD1^G93A^ mice and did not modify the score in the beam balance test (Figure 4A-D).

**Figure 4.**
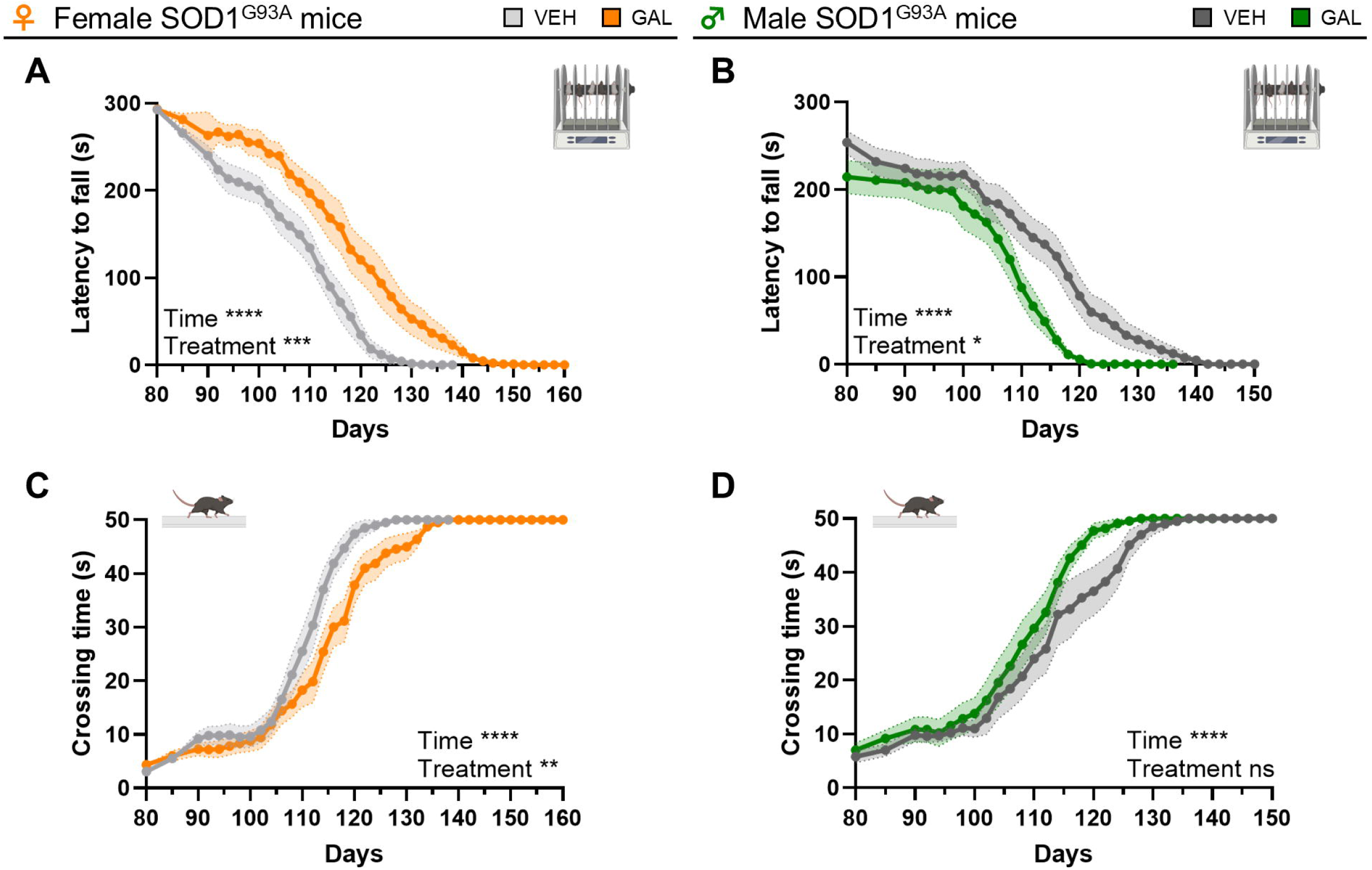
Motor coordination in GAL and vehicle treated SOD1^G93A^ mice. (**A-B**) Quantification of the time spent on the rotarod of female (A) and male (B) SOD1^G93A^ mice treated with vehicle (VEH) or 10 mg/kg/day GAL (n=10-12). (**C-D**) Quantification of beam crossing time of female (C) and male (D) SOD1^G93A^ mice treated with VEH or 10 mg/kg/day GAL (n=10-12). * p<0.05, ** p<0.01, *** p<0.001, **** p<0.0001, mixed-effects model with Geisser-Greenhouse correction for repeated measures.

The analysis of motor skills unveiled a significant amelioration of both extension reflex and gait scores in GAL-treated SOD1^G93A^ females compared to vehicle, while opposite modifications were found in males (Figure 5A-D).

**Figure 5.**
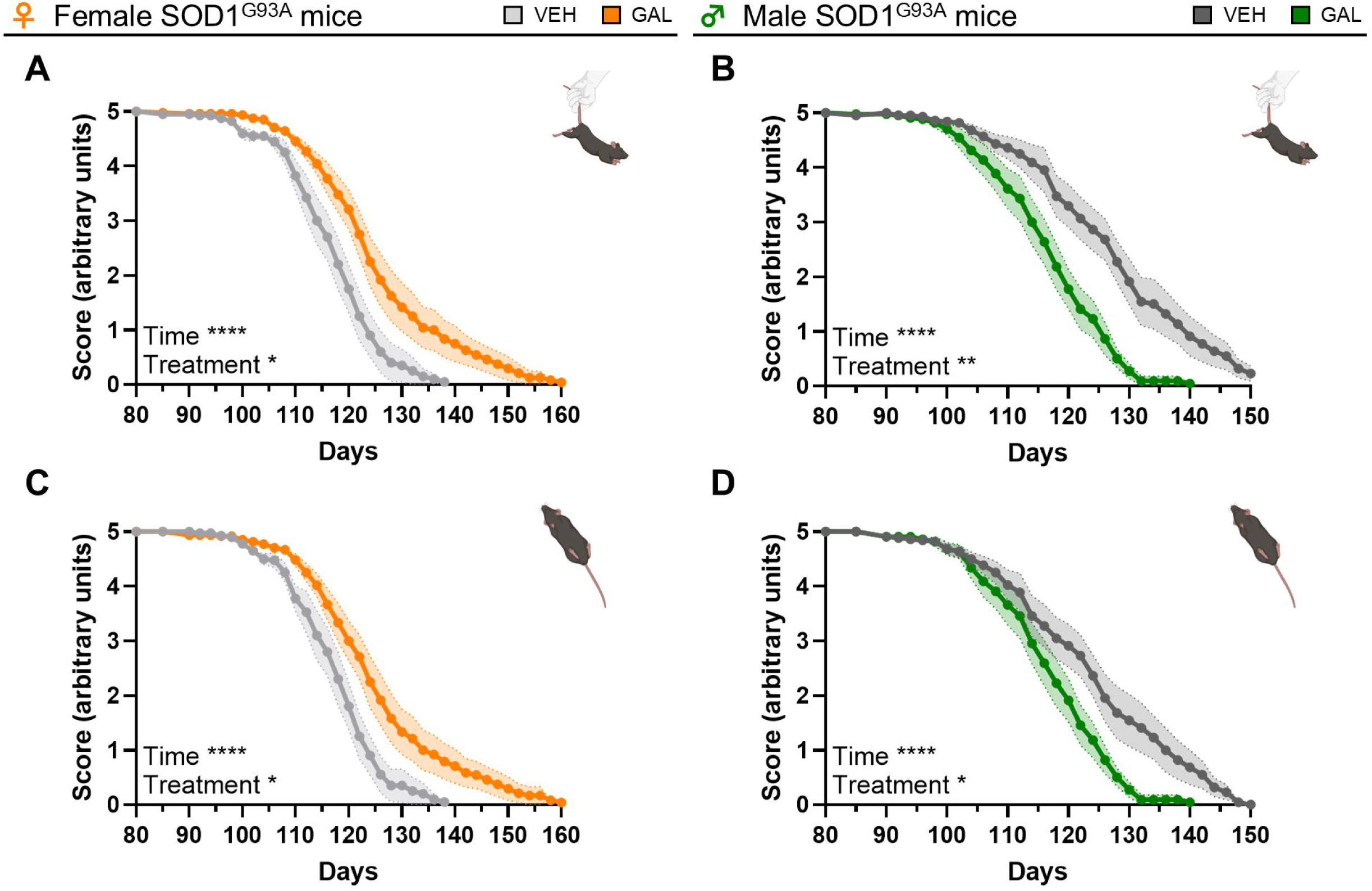
Motor skills in GAL and vehicle treated SOD1^G93A^ mice. (**A-B**) Quantification of the hindlimb extension reflex of female (A) and male (B) SOD1^G93A^ mice treated with vehicle (VEH) or 10 mg/kg/day GAL (n=10-12). (**C-D**) Quantification of the gait score of female (C) and male (D) SOD1^G93A^ mice treated with VEH or 10 mg/kg/day GAL (n=10-12). * p<0.05, ** p<0.01, **** p<0.0001, mixed-effects model with Geisser-Greenhouse correction for repeated measures.

Finally, treatment with GAL significantly improved forelimb grip strength in SOD1^G93A^ females but not in males, while it did not promote any amelioration of the hanging wire test score in either sex (Figure 6A-D).

**Figure 6.**
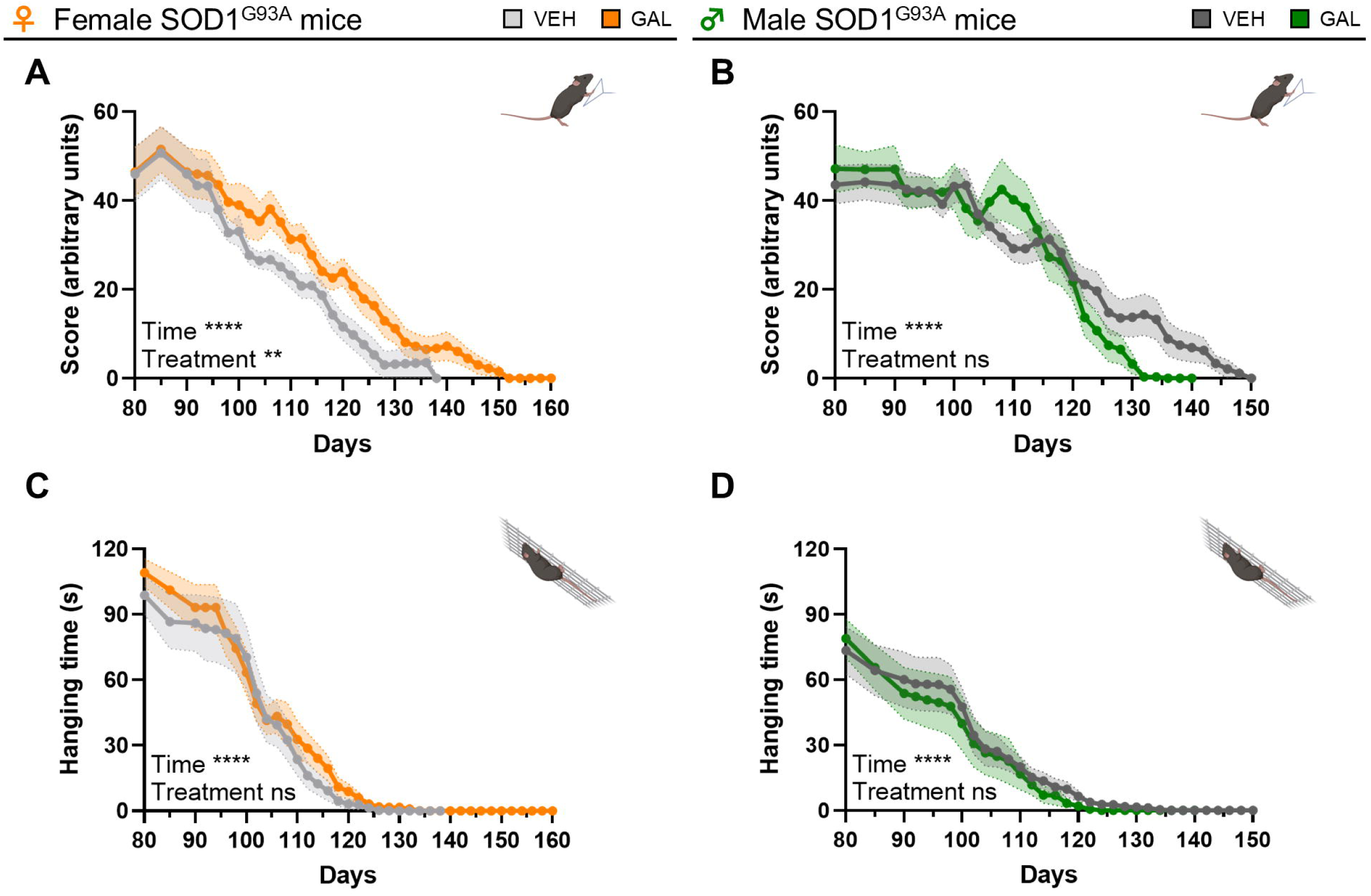
Muscle strength in GAL and vehicle treated SOD1^G93A^ mice. (**A-B**) Quantification of the forelimb grip strength score of female (A) and male (B) SOD1^G93A^ mice treated with vehicle (VEH) or 10 mg/kg/day GAL (n=10-12). (**C-D**) Quantification of the hindlimb hanging wire test duration of female (C) and male (D) SOD1^G93A^ mice treated with VEH or 10 mg/kg/day GAL (n=10-12). ** p<0.01, **** p<0.0001, mixed-effects model with Geisser-Greenhouse correction for repeated measures.

Altogether, these data indicate that GAL treatment significantly improves motor coordination, motor skills, and muscle strength in female SOD1^G93A^ mice, while no consistent effects are observed in males.

### 3.5. Chronic *in vivo* treatment with a selective GPR17 agonist counteracts pathological GPR17 upregulation and restores oligodendrocyte maturation in the spinal cord of female SOD1^G93A^ mice

To investigate whether the beneficial effects of GAL on survival and disease progression of female SOD1^G93A^ mice were causally associated with changes in the oligodendroglial populations, we performed immunohistochemical analyses at the late symptomatic stage (P125). Given the prominent vulnerability of lower motor neuron circuits in this model, analyses were focused on the ventral gray matter of lumbar spinal cord, a region preferentially affected in SOD1^G93A^ mice [35].

In female mice, chronic GAL treatment induced a marked shift in the oligodendroglial lineage, indicative of enhanced maturation (Figure 7). Specifically, GAL significantly reduced the density of NG2 early OPCs and GPR17 immature oligodendrocytes compared to vehicle, while concomitantly increasing the number of ASPA mature oligodendrocytes in the ventral gray matter (Figure 7A-). In contrast, in male SOD1^G93A^ mice, GAL treatment led to a reduction in NG2 OPCs but did not significantly affect the abundance of GPR17 immature cells or ASPA mature oligodendrocytes (Figure 7F-I). Moreover, GAL treatment did not alter the total number of OLIG2 oligodendrocyte lineage cells in either female or male SOD1^G93A^ mice (Figure 7E, J), suggesting that the changes in NG2^+^, GPR17 and ASPA cells observed in females result from a redistribution of maturation states within the oligodendrocyte lineage rather than changes in cell number due to proliferation or cell death.

**Figure 7.**
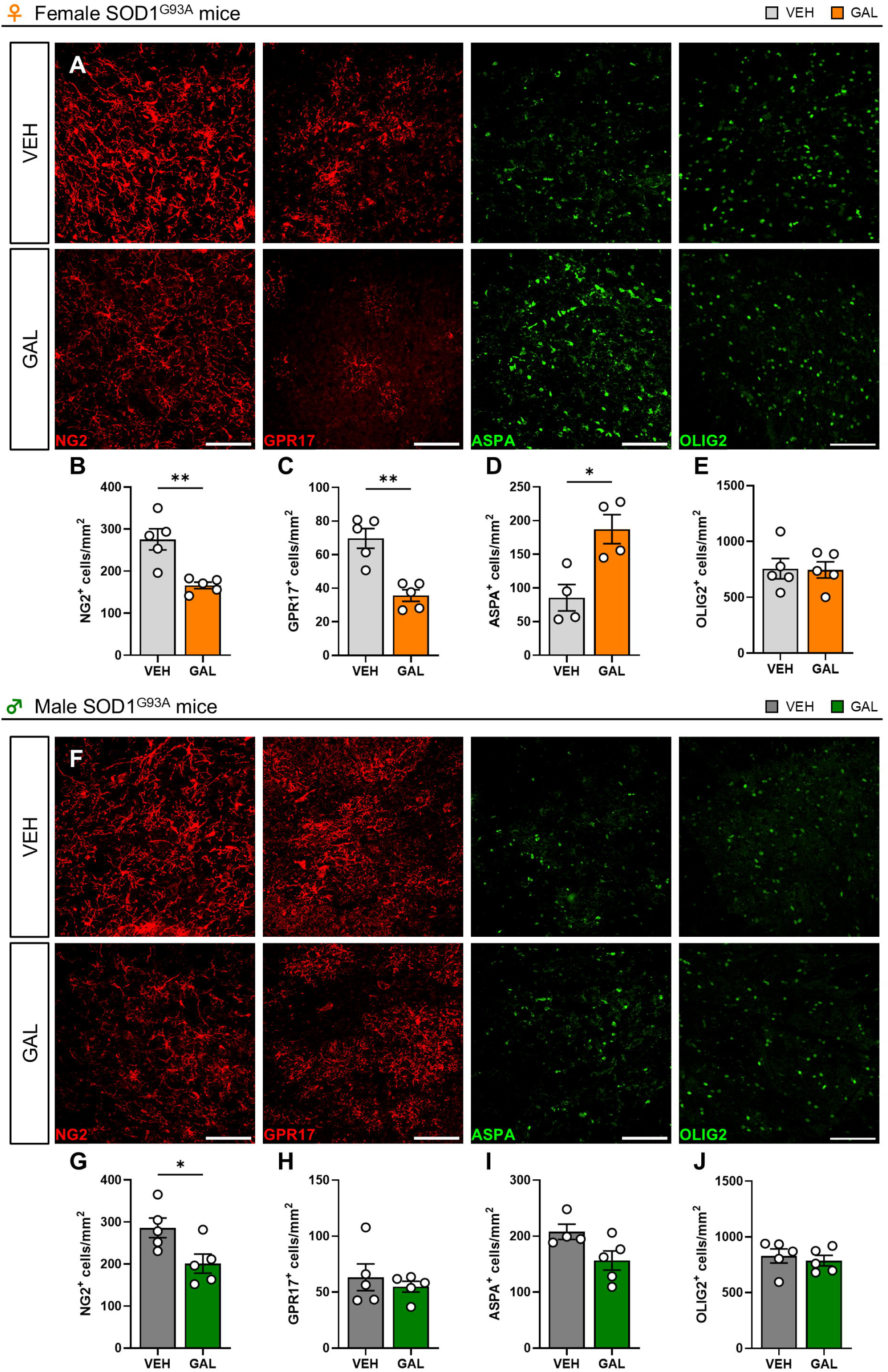
Oligodendrocyte maturation in the spinal cord of GAL and vehicle treated SOD1^G93A^ mice. (**A**) Representative images of cells stained for NG2, GPR17, ASPA, and OLIG2 in the ventral gray matter of the lumbar spinal cord of P125 female SOD1^G93A^ mice treated with 10 mg/kg/day GAL or vehicle (VEH). Nuclei were labelled with Hoechst33258. Scale bar: 100 µm. (**B-E**) Quantification of the density of cells expressing NG2 (B), GPR17 (C), ASPA (D), and OLIG2 (E) in the ventral gray matter of the lumbar spinal cord of P125 female SOD1^G93A^ mice treated with 10 mg/kg/day GAL or VEH (n=4-5). * p<0.05, ** p<0.01, Student’s t-test. (**F**) Representative images of cells stained for NG2, GPR17, ASPA, and OLIG2 in the ventral gray matter of the lumbar spinal cord of P125 male SOD1^G93A^ mice treated with 10 mg/kg/day GAL or VEH. Nuclei were labelled with Hoechst33258. Scale bar: 100 µm. (**G-J**) Quantification of the density of cells expressing NG2 (G), GPR17 (H), ASPA (I), and OLIG2 (J) in the ventral gray matter of the lumbar spinal cord of P125 male SOD1^G93A^ mice treated with 10 mg/kg/day GAL or VEH (n=4-5). * p<0.05, Student’s t-test.

These findings indicate that GAL promotes oligodendroglial differentiation *in vivo* in a sex-dependent manner, counteracting pathological GPR17 upregulation and favoring transition of blocked immature precursors towards mature oligodendrocytes in female SOD1^G93A^ mice.

### 3.6. Chronic *in vivo* treatment with a selective GPR17 agonist improves myelin integrity and motor neuron survival in the spinal cord of female SOD1^G93A^ mice

Given the GAL-induced beneficial effects on oligodendroglial maturation observed in female SOD1^G93A^ mice, we next assessed whether these cellular changes translated into improved myelin integrity and motor neuron preservation *in vivo*.

In female SOD1^G93A^ mice, chronic GAL treatment resulted in a significant increase in MBP area fraction, indicating enhanced myelin content (Figure 8A-B). Consistently, quantitative co-localization analyses of MBP and NF-H revealed a significant increase in myelinated axons in GAL-treated female mice, supporting improved myelin–axon coupling (Figure 8A-C). These findings were further corroborated by spectral confocal reflectance (SCoRe) microscopy, a label-free imaging technique utilized to visualize intact myelin in the CNS based on the high refractive index of compacted myelin compared with the surrounding tissue [41,42]. This approach showed a significant increase in the compact myelin–positive (SCoRe ) area in GAL-treated females compared with vehicle-treated controls (Figure 8A-D). Importantly, these improvements in myelin integrity were associated with a significant preservation of Hb9 spinal motor neurons, indicating protection of vulnerable neuronal populations (Figure 8A-E). In contrast, male SOD1^G93A^ mice did not exhibit significant GAL-induced changes in MBP area fraction or MBP/NF-H co-localization, while a trend toward increased SCoRe signal could be observed. Similarly, motor neuron survival was not affected by GAL in males (Figure 8F-J).

**Figure 8.**
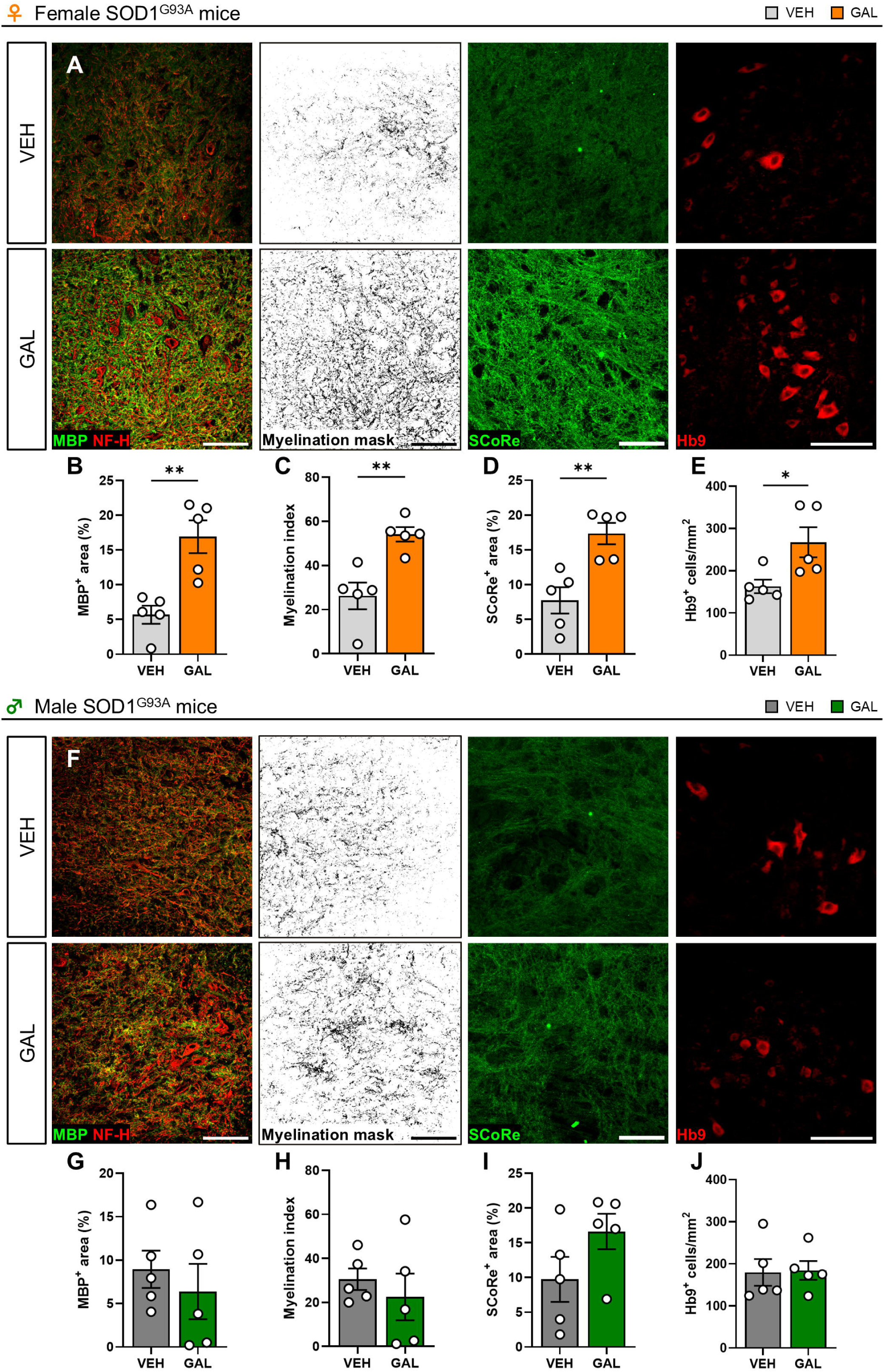
Myelin integrity and motor neuron survival in the spinal cord of GAL and vehicle treated SOD1^G93A^ mice. (**A**) Representative images of MBP and NF-H double staining, MBP/NF-H co-localization mask, SCoRe^+^ compact myelin signal, and Hb9^+^ motor neurons in the ventral gray matter of the lumbar spinal cord of P125 female SOD1^G93A^ mice treated with 10 mg/kg/day GAL or vehicle (VEH). Scale bar: 100 µm. (**B-E**) Quantification of MBP^+^ area fraction (B), myelination index (C), SCoRe^+^ area fraction (D), and Hb9^+^ cell density (E) in the ventral gray matter of the lumbar spinal cord of P125 female SOD1^G93A^ mice treated with 10 mg/kg/day GAL or VEH (n=5). * p<0.05, ** p<0.01; Student’s t-test. (**F**) Representative images of MBP and NF-H double staining, MBP/NF-H co-localization mask, SCoRe^+^ compact myelin signal, and Hb9^+^ motor neurons in the ventral gray matter of the lumbar spinal cord of P125 male SOD1^G93A^ mice treated with 10 mg/kg/day GAL or VEH. Scale bar: 100 µm. (**G-J**) Quantification of MBP^+^ area fraction (G), myelination index (H), SCoRe^+^ area fraction (I), and Hb9^+^ cell density (J) in the ventral gray matter of the lumbar spinal cord of P125 male SOD1^G93A^ mice treated with 10 mg/kg/day GAL or VEH (n=5).

These data demonstrate that GAL treatment fosters myelin repair and supports motor neuron survival in SOD1^G93A^ mice in a sex-dependent manner.

### 3.7. Chronic *in vivo* treatment with a selective GPR17 agonist counteracts reactive gliosis in the spinal cord of SOD1^G93A^ mice

Finally, to assess whether the beneficial effects of chronic GAL administration extended to neuroinflammatory responses associated with ALS progression, we analyzed markers of reactive gliosis in the lumbar spinal cord of SOD1^G93A^ mice. Given that GPR17 expression is restricted to the oligodendroglial lineage, any positive effects on microglia or astrocytes would be expected to occur indirectly, potentially because of improved myelin integrity and motor neuron preservation.

In female SOD1^G93A^ mice, GAL treatment resulted in a significant reduction in the Iba1 microglia area fraction (Figure 9A-B), together with a marked attenuation of GFAP astrocytic reactivity (Figure 9A-C). In parallel, GAL significantly reduced SERPINA3N immunoreactivity (Figure 9A-D), indicating a dampening of disease-associated glial states [26,44]. In male mice, GAL induced a not-significant downward trend in the Iba1 microglial/macrophage area (Figure 9E-F), while GFAP and SERPINA3N area fractions were significantly reduced (Figure 9E-H).

**Figure 9.**
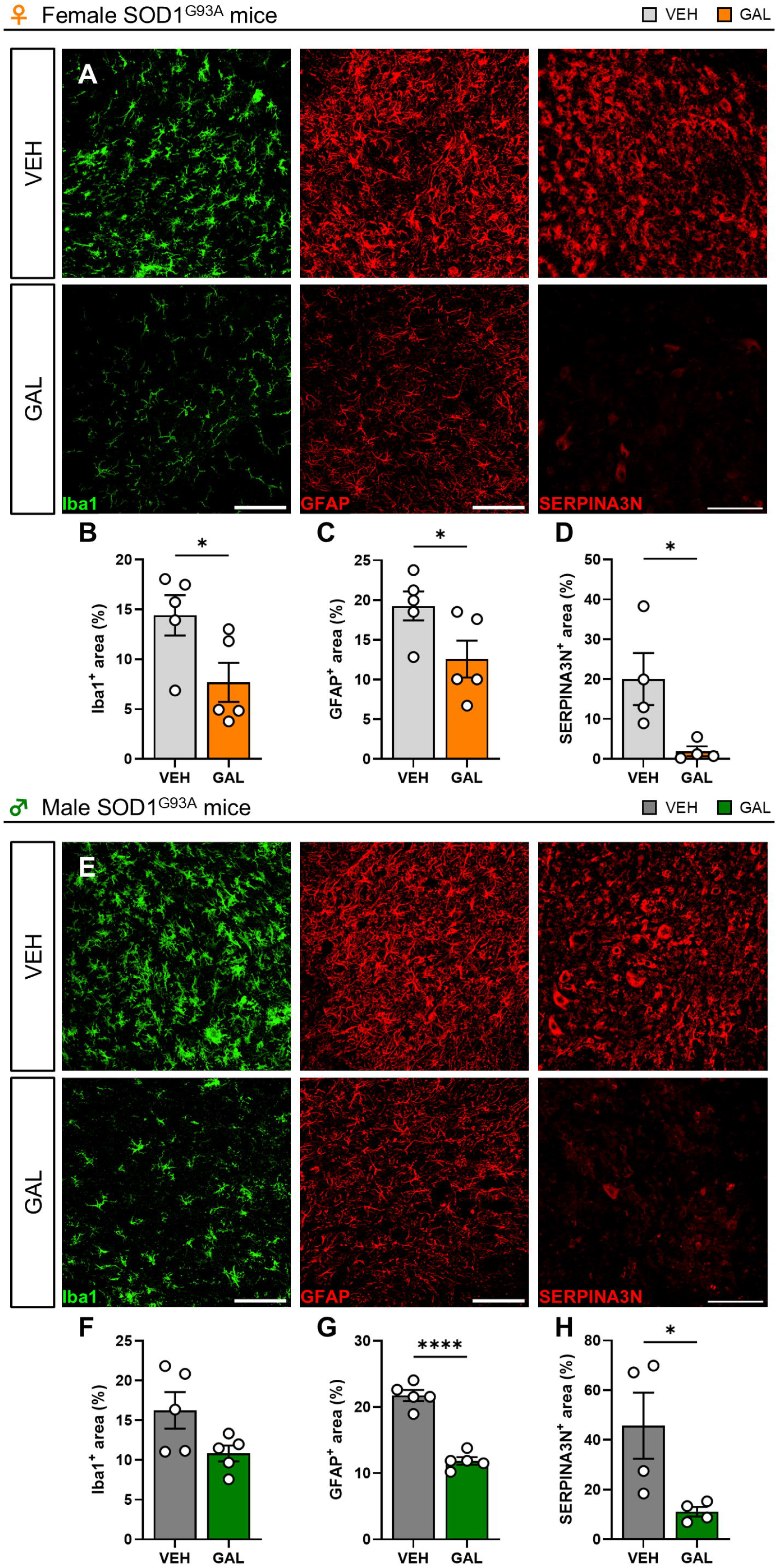
Reactive gliosis in the spinal cord of GAL and vehicle treated SOD1^G93A^ mice. (**A**) Representative images of Iba1^+^ microglia/macrophages, GFAP^+^ astrocytes, and SERPINA3N^+^ disease-associated glia in the ventral gray matter of the lumbar spinal cord of P125 female SOD1^G93A^ mice treated with 10 mg/kg/day GAL or vehicle (VEH). Scale bar: 100 µm. (**B-D**) Quantification of Iba1^+^ area fraction (B), GFAP^+^ area fraction (C), and SERPINA3N^+^ area fraction (D) in the ventral gray matter of the lumbar spinal cord of P125 female SOD1^G93A^ mice treated with 10 mg/kg/day GAL or VEH (n=4-5). * p<0.05, Student’s t-test. (**E**) Representative images of Iba1^+^ microglia/macrophages, GFAP^+^ astrocytes, and SERPINA3N^+^ disease-associated glia in the ventral gray matter of the lumbar spinal cord of P125 male SOD1^G93A^ mice treated with 10 mg/kg/day GAL or VEH. Scale bar: 100 µm. (**F-H**) Quantification of Iba1^+^ area fraction (F), GFAP^+^ area fraction (G), and SERPINA3N^+^ area fraction (H) in the ventral gray matter of the lumbar spinal cord of P125 male SOD1^G93A^ mice treated with 10 mg/kg/day GAL or VEH (n=4-5). * p<0.05, **** p<0.0001, Student’s t-test.

Taken together, these data indicate that chronic treatment with GAL blunts reactive gliosis in the spinal cord of SOD1^G93A^ mice, with more robust effects in females. Although these analyses do not establish the mechanism underlying the glial changes induced by GAL, they identify a potentially relevant effect on the spinal cord inflammatory environment that warrants dedicated investigation in future studies.

## 4. Discussion

In the present study, we identify the oligodendroglial GPR17 receptor as a conserved and pharmacologically actionable target in human ALS and demonstrate that its modulation with a selective agonist can induce robust, sex-dependent therapeutic effects in a widely used preclinical ALS model. By integrating human post-mortem tissues with mechanistic in vitro analyses and chronic in vivo pharmacological intervention, our findings suggest that GPR17 upregulation marks a pathological oligodendroglial state that can be therapeutically redirected to restore myelin integrity and improve disease outcome in ALS.

A key translational advance of this study is the demonstration that GPR17 dysregulation identified in ALS mouse models is conserved in human disease. Extending our previous observations of elevated GPR17 protein levels and increased density of GPR17 cells in the spinal cord of SOD1G93A mice [9,20], here we show that GPR17 is consistently upregulated in the human ALS spinal cord, as evidenced by concordant findings from bulk and single-nucleus RNA-sequencing analyses and corroborated by immunohistochemistry. Importantly, the human cohorts analyzed include both sporadic ALS cases and patients carrying distinct genetic mutations, indicating that GPR17 upregulation is not confined to SOD1-linked ALS but instead represents a shared pathological feature across multiple ALS subtypes. Although the available histological cohort was not sufficiently powered for sex-stratified analyses, the comparable GPR17 cell densities observed in male and female donors, together with transcriptomic datasets comprising balanced numbers of male and female subjects, support the association between GPR17 upregulation and ALS pathology. Nevertheless, larger sex-balanced cohorts will be needed to formally determine any potential sex-dependent modulation of GPR17 dysregulation in human ALS. At the cellular level, GPR17 upregulation was found to be highly specific to BCAS1-expressing immature oligodendrocytes, with no detectable expression in mature oligodendrocytes, astrocytes, microglia, or neurons. This cellular specificity is consistent with our previous observations in post-mortem tissue from human cases with multiple sclerosis [45] and ischemic stroke [27] and supports a targeted alteration of the oligodendroglial lineage. Such lineage restriction confirms the specificity of the pharmacological effects observed and suggests that therapeutic modulation of GPR17 may limit off-target effects on other neural cell types. The convergence of molecular and histological evidence across species, disease etiologies and analytical platforms reinforces the translational relevance of GPR17 and strengthens its potential as a disease-relevant, oligodendrocyte-specific therapeutic target in human ALS.

One potential concern in targeting GPR17 with agonists is the apparent paradox of pharmacologically engaging a receptor that is already pathologically upregulated in disease. To date, the most intuitive and widely pursued strategy for GPR17 targeting has been the use of antagonists, aimed at blocking excessive signaling [20,46–51]. However, in the case of GPR17, such an approach may carry important limitations, as receptor activation is required during the early phases of oligodendrocyte lineage progression [12] and indiscriminate antagonism could interfere with its physiological functions at stages (or CNS sites) where GPR17 signaling is still necessary. In contrast, agonist-based strategies offer a mechanistically distinct and potentially more physiological mode of intervention in chronic disease settings. Sustained agonist-mediated stimulation of GPR17 does not result in persistent receptor activation; rather, it triggers canonical GPCR regulatory mechanisms [52] including desensitization, internalization and subsequent downregulation, thereby relieving the differentiation block imposed by sustained receptor overexpression [16,21,22]. Importantly, this process closely recapitulates the physiological dynamics of GPR17 signaling during oligodendrocyte maturation, which is tightly regulated by extracellular cues and requires timely receptor downregulation to enable progression toward a mature, myelinating phenotype [16]. Fueled by this hypothesis, our previous work identified a class of novel and selective GPR17 agonists [24,43], whose lead-compound ASN was shown to concentration-dependently inhibit forskolin-stimulated cAMP accumulation in primary OPCs, consistent with activation of the canonical G_i_ signaling pathway downstream of GPR17 [23]. Here, our data show that treatment with ASN was able to induce redistribution of the receptor from the plasma membrane toward intracellular vesicular compartments *in vitro* and to enhance terminal differentiation and release of neurotrophic factors in SOD1^G93A^ OPCs. Although these assays do not provide a quantitative analysis of receptor trafficking kinetics, our findings are consistent with agonist-induced receptor internalization and support the idea that sustained and continuous GPR17 engagement may promote functional normalization of receptor activity rather than exacerbation of pathological signaling. This is consistent with our recent *in vitro* data showing that GAL treatment is able to rescue oligodendrocyte maturation and myelination under disease-related inflammatory conditions [53]. *In vivo*, chronic GAL treatment reduced the number of immature GPR17 cells while increasing mature oligodendrocytes, a pattern consistent with an attenuation of pathological GPR17 signaling. However, it is important to note that these data do not directly demonstrate agonist-induced receptor desensitization, internalization, or downregulation in spinal cord oligodendrocytes *in vivo.* Nevertheless, the possibility that prolonged GAL exposure functionally dampens GPR17 signaling remains biologically plausible, as analogous pharmacological paradigms have proven successful for other GPCRs in chronic pathological contexts, where prolonged agonist exposure is indeed exploited to induce receptor downregulation and therapeutic benefit. This is the case, for instance, of gonadotrophin releasing hormone (GnRH) agonists in endocrine pathologies and cancer, where continuous stimulation paradoxically suppresses receptor signaling via internalization and downregulation [54]. A closely related example in the neuroglial field is fingolimod (FTY720), which has been proven to be clinically effective in multiple sclerosis by modulating immune cell trafficking and by exerting direct pro-myelinating and neuroprotective effects in the CNS [55,56]. Interestingly, fingolimod has been shown to act as a functional antagonist of sphingosine-1-phosphate receptors, which initially acts as an agonist but induces receptor internalization and degradation upon sustained exposure [57]. Recent evidence shows that GPR17 may display constitutive activity [58], suggesting that receptor signaling could occur even in the absence of endogenous ligands. Although the present study did not specifically address basal receptor activity, this observation is compatible with our proposed mechanism. In disease states characterized by sustained GPR17 overexpression, increased constitutive activity may contribute to the persistence of an immature oligodendroglial phenotype, whereas chronic agonist exposure would be expected to promote receptor desensitization and internalization, thereby reducing functional receptor signaling and relieving the differentiation block. Together, these considerations indicate that agonist-driven modulation of GPR17 represents a rational, well-validated, and disease-informed strategy harnessing endogenous GPCR regulatory mechanisms to restore oligodendroglial differentiation.

Chronic in vivo administration of the optimized GPR17 agonist GAL [24] translated the cellular and molecular effects observed *in vitro* into meaningful functional benefits, albeit in a strikingly sex-dependent manner. In female SOD1^G93A^ mice, GAL significantly extended survival, delayed body weight loss, and improved multiple motor outcomes while promoting oligodendrocyte maturation, fostering remyelination, and protecting spinal motor neurons. Of note, GAL also increased food intake in female, but not male, SOD1^G93A^ mice, consistent with its effects on body weight. These data are also in line with previous studies showing sex-dependent metabolic phenotypes in GPR17 genetic models, in which GPR17 signaling was linked to energy balance, feeding behavior and metabolic regulation in a sex- and context-dependent manner [59,60]. However, it remains unclear whether this contributes to the therapeutic benefit, possibly through a direct role of oligodendroglial GPR17 in regulating feeding behavior [61] or, conversely, reflects improved feeding efficiency resulting from the overall amelioration of disease progression. Importantly, comparable sex-dependent beneficial effects were obtained using the non-selective GPR17 antagonist montelukast [20], indicating that, despite using different molecular mechanisms, both agonist- and antagonist-based modulation of GPR17 converge on similar disease-modifying outcomes in females. The reproducibility of these effects across different pharmacological classes, each with distinct mechanisms of receptor engagement, strongly supports *in vivo* target engagement of GPR17 by both montelukast and GAL. It also reinforces the notion that GPR17 represents a disease-relevant target in ALS, whose modulation yields consistent benefits in females. In contrast, GAL treatment failed to confer benefit in male SOD1^G93A^ mice and, in some behavioral paradigms, was even associated with worsened performance, raising important mechanistic and translational considerations. Such sex-specific therapeutic responses may reflect intrinsic biological differences in pathophysiological processes, including dimorphisms of glial biology, neuroinflammatory responses, and disease progression dynamics. Sex-dependent differences in glial reactivity [62], (re)myelination capacity [63,64], and GPCR regulation [65] are increasingly recognized and are directly relevant to GPR17 signaling. Sex hormones may be particularly relevant in this context, as estrogenic signaling has been implicated in oligodendrocyte differentiation, lipid metabolism, myelin repair and neuroprotection, processes that are closely related to the cellular effects of GPR17 modulation. Moreover, sex-dependent differences in GPCR expression, downstream coupling, receptor trafficking and desensitization may influence the pharmacodynamic response to chronic GPR17 agonism. Although, to our knowledge, no definitive evidence currently demonstrates sex-specific differences in GPR17 expression or signaling in ALS patients, broader sex-related differences in ALS incidence, clinical phenotype, progression and molecular signatures are well recognized and may influence therapeutic responsiveness. Accordingly, female SOD1^G93A^ mice display slower disease progression and better motor performance than males [66,67], mirroring the reduced susceptibility and slower progression observed in women with ALS [68,69]. Consequently, initiating treatment at the same postnatal time point may engage distinct disease stages in males and females, potentially accounting for divergent therapeutic outcomes. This may indicate that the therapeutic window in male mice is already closed by the time GAL treatment is started. Considering that sex is an independent modifier of ALS incidence, progression and therapeutic response, our findings underscore the need for precision pharmacology and gender-informed therapeutic strategies. However, the mechanisms underlying these sex differences in ALS, including the possible interaction between sex hormones, GPR17 signaling and oligodendroglial repair capacity, remain largely unexplored [70,71] and warrant further investigation.

Importantly, GAL has been previously characterized as a selective GPR17 agonist [24], and our data confirm that GPR17 expression in the CNS is primarily restricted to cells of the oligodendroglial lineage. In this context, the reduction of microglial and astrocytic reactivity observed following GAL treatment supports a predominantly non–cell-autonomous mechanism of action rather than a direct effect of GAL on these cell populations. In females, these changes are coherent with the broader histological and functional improvement induced by GAL, and may therefore arise, at least in part, as secondary consequences of enhanced oligodendrocyte maturation, improved myelin integrity, and motor neuron preservation. Both microglia and astrocytes are highly sensitive to myelin disruption and debris, which act as potent pro-inflammatory stimuli through pattern-recognition receptors, lipid-sensing pathways and inflammasome activation [72,73]. Accumulation of myelin debris is known to sustain chronic microglial activation, impair phagocytic resolution and promote astrocytic reactive states, thereby amplifying neuroinflammation [39,74–76]. In this context, the significant remyelination observed in GAL-treated females is likely to contribute to the reduction of myelin-derived danger signals and, consequently, to the dampening of glial reactivity. In males, however, the attenuation of selected gliosis markers occurred despite limited effects on oligodendrocytes, myelin, or motor neurons, indicating that these reactive changes cannot be simply interpreted as a downstream readout of structural rescue and suggesting a more complex interplay between oligodendrocytes and surrounding glial cells. Beyond their role in myelination, oligodendrocytes are increasingly recognized as active contributors to CNS immune homeostasis, with the capacity to influence glial responses through metabolic support, lipid handling, cytokine signaling, and regulation of the extracellular milieu [44,77]. Accordingly, the reduction of SERPINA3N-positive disease-associated glial states following GAL treatment further supports the existence of functionally relevant crosstalk between oligodendrocytes, astrocytes, and microglia. Although deep investigation of these interactions goes beyond the scope of this study, these data represent an important observation that is consistent with the multicellular and non–cell-autonomous nature of ALS pathology. Future studies specifically designed to dissect the reciprocal interactions between oligodendrocytes, astrocytes, and microglia will be necessary to determine how GPR17 modulation in oligodendrocytes contributes to reshaping the inflammatory microenvironment. Moreover, since oligodendroglial dysfunction emerges early in disease progression, before overt neuroinflammation and neurodegeneration are evident [4,5,9,78], these findings support the idea that targeting oligodendrocytes via GPR17 may influence broader disease-relevant cellular interactions in ALS. Overall, our data identify GPR17 modulation as a promising oligodendrocyte-centered strategy, while also pointing to glial crosstalk as an important area for future investigation.

As with all preclinical studies, limitations should be acknowledged. The SOD1^G93A^ mouse model represents only a subset of ALS cases and does not fully recapitulate the marked genetic, molecular and clinical heterogeneity of the human disease [79]. Nevertheless, this limitation is substantially mitigated by our data showing that GPR17 dysregulation is conserved in human ALS spinal cord, across cohorts comprising both sporadic cases and multiple genetic backgrounds, thereby supporting the broader disease relevance of this pathway beyond SOD1-linked ALS. We acknowledge that ultrastructural analyses such as transmission electron microscopy with g-ratio assessment would be needed to conclusively define the extent of myelin repair; however, the combined MBP, MBP/NF-H co-localization, and SCoRe analyses reported here consistently support GAL ability to promote myelin integrity. Finally, the present study evaluated a single dosing regimen of GAL, selected based on the available pharmacokinetic data supporting brain penetration compatible with its activity [24], initiated at the early symptomatic stage to model a therapeutic rather than a preventive intervention. While this design strengthens translational relevance, future studies will be required to systematically define optimal dosing, treatment duration, and therapeutic windows, as well as assess long-term safety. These efforts will be particularly important considering the pronounced sex-dependent effects observed, which underscore the need to refine GPR17-targeted strategies in a sex-informed manner rather than assuming uniform pharmacological responses. Collectively, these considerations provide a clear roadmap for the next phase of preclinical development, aimed at advancing GPR17 modulation as a precision, oligodendrocyte-targeted therapeutic strategy in ALS.

In conclusion, our work identifies the oligodendroglial GPR17 receptor as a conserved player in ALS pathology and supports GPR17 desensitization with selective agonists as a viable strategy to restore oligodendrocyte maturation, enhance remyelination, and slow disease progression. The data obtained with GAL represent a proof-of-concept for the efficacy of a therapeutic approach based on the *in vivo* administration of GPR17 agonists. By providing human validation, mechanistic insight, and *in vivo* efficacy, these findings support the further development of GPR17-targeted compounds for ALS treatment.

## Acknowledgements

We acknowledge Elisabetta Bonfanti (Università degli Studi di Milano), Giuseppe Marazzotta (Università degli Studi di Genova), and Ulla Damgaard Munk (University of Southern Denmark) for skilled technical assistance. This work was supported by AriSLA, (grants GPR17ALS to MF; GPR17ALS-1 and DORALS to MF and TB) and by TargetALS (grant BSP-TAL006 to MF). The research was also supported by PRIN-Progetti di Ricerca di Interesse Nazionale (grant 2017NSXP8J to MPA) and Fondazione Italiana Sclerosi Multipla (grant 2017/R/1 to MPA). The authors are also grateful to the Foundation Bellandi Bernardoni (Genoa) for supporting this research, and to the master’s students Mirko Nitro, Debora Legnaro, and Martina Scaglione for their valuable support. Part of this work was carried out at NOLIMITS, an advanced imaging facility established by Università degli Studi di Milano. CISUP is acknowledged for access to the microscopy facility of University of Pisa. Cartoons have been created with BioRender.com.

## Data statement

Further information and data that support the findings of this study are available from the corresponding author upon reasonable request.

